# Mapping the AI Life Sciences Landscape in Greece: A National Survey and Bibliometric Comparison with Global Trends

**DOI:** 10.1101/2025.11.19.689212

**Authors:** Eleni Adamidi, Serafeim Chatzopoulos, Alexandros C. Dimopoulos, Anastasia Krithara, Anastasios Nentidis, Nikos Pechlivanis, Fotis Psomopoulos, Thanasis Vergoulis, Kleanthis Vichos

## Abstract

Artificial intelligence is increasingly used in Life Sciences, though the pace and direction of adoption varies widely across countries. To map the Greek landscape, we combined two complementary approaches, a data-driven analysis of 916,824 AI-related life-science papers harvested from OpenAlex and PubMed and a targeted short survey at a national level aimed towards researchers, engineers, and support staff to be used as supporting material. We tagged each publication with Medical Subject Headings (MeSH) and compared topic frequencies between articles linked to at least one Greek institution and the rest of the world. Greek-affiliated outputs are disproportionately concentrated under the theme of methodology and algorithm-development, whereas the global corpus is dominated by disease-focused, organism-centered and clinical applications. Statistical contrasts across three MeSH hierarchy levels exposed clear national strengths in machine learning techniques and analytical tools, alongside under-representation in translational, patient-centred research. Survey responses reinforce these patterns: participants highlight limited access to well-curated biomedical data, constrained computational resources and difficulty recruiting cross-disciplinary talent as the chief barriers to progress. They advocate community-building actions (e.g., hackathons, postgraduate training, shared data infrastructures) that could realign national efforts with international practice while capitalizing on existing expertise in core-methodology development. Overall this study combines bibliometric evidence with community perspectives and provides a comprehensive overview of AI activity in Life Sciences in Greece, highlighting potential thematic strengths and gaps.

## Introduction

Artificial Intelligence (AI) has rapidly evolved over the past decade, becoming one of the most influential technological advancements across scientific fields. The impact is particularly evident in research and innovation, where it has reshaped methodologies, enabled new discoveries, and offered solutions to previously complex problems. AI methods are increasingly used in Life Sciences to assist the analysis of biological data facilitating the extraction of valuable insights and the discovery of latent patterns. Their applications span a wide spectrum, including predictive modeling in genomics and systems biology, image analysis for diagnostics, optimization of experimental workflows, and, more recently, the use of Large Language Models (LLMs) for clinical data interpretation.

As the integration of AI in Life Sciences becomes increasingly widespread, a growing number of international initiatives have emerged to explore the infrastructural, ethical, and research implications of this transformation, both across and within individual communities or countries. One such effort is the Report by UKRI [1] on digital research infrastructure requirements for AI, which provides a national-level perspective on the challenges and opportunities associated with AI adoption across disciplines. It identifies critical needs in terms of skills, data management, compute infrastructure, and interdisciplinary collaboration. Inspired by the scope of this UK-based initiative, we sought to carry out a similar effort tailored to the context of Greece.

Greece has a growing research ecosystem in Life Sciences and computational technologies, yet there is still limited systematic understanding of how AI is currently being applied in this domain at a national level. Our study aims to fill this gap by mapping the intersection between AI and Life Sciences in Greece. This mapping is timely, as recent investments in national research infrastructure and the emergence of collaborative initiatives (e.g., ELIXIR-GR) have begun to support computationally intensive work in biology, biomedicine, and related fields.

To investigate the state of AI activity in Life Sciences in Greece, we adopted a twofold approach. First, we conducted a national survey directed at researchers, engineers, support staff, and institutional representatives engaged in AI and Life Sciences research. The survey aimed to capture perceptions regarding current capabilities, infrastructure, key research areas, skill gaps, and future needs. Second, we carried out a data-driven analysis using bibliographic datasets from OpenAlex [2] and PubMed [3]. This involved retrieving a large collection of publications at the intersection of Life Sciences and AI-related fields (e.g., machine learning, natural language processing) and enriching them with valuable metadata, including institutional affiliations, topics (OpenAlex concepts), and subject headings (MeSH terms).

The combination of survey responses and bibliometric analysis enabled us to capture both perspectives from the Greek community and evidence of research activity in the AI–Life Sciences interface. Our findings aim to offer insights into thematic trends, institutional expertise, and training landscapes in Greece, and to highlight how they compare with broader international patterns. Moreover, the analysis supports a discussion of national research priorities and infrastructural investments needed to facilitate further AI integration in the Life Sciences.

## Results

The overarching analysis process consisted of consecutive data retrieval, annotation and filtering steps, followed by a comparative statistical analysis between the selected groups. OpenAlex served as the primary source of bibliographic metadata. By linking each publication’s DOI to its corresponding MeSH in PubMed, we compiled a richly annotated corpus of articles positioned at the intersection of Medicine and Biology, and further associated with topics such as Natural Language Processing, Artificial Intelligence, or Machine Learning (see Methods). The final dataset was then divided into two groups, “Greece” and “World”, depending on whether a publication listed at least one institution located in Greece among its affiliations. This division allowed us to explore how the thematic focus of AI–Life Sciences research in Greece aligns with – or diverges from – the global landscape.

### Comparative overview of Greek and global AI–Life Sciences research

To understand the thematic structure of AI Life Sciences research, we focused on the main topics of these publications, based on their Medical Subject Headings (MeSH) [4] annotations. The MeSH thesaurus is a controlled vocabulary developed and maintained by the National Library of Medicine (NLM). The basic entity in MeSH is the MeSH descriptor, which is a set of related terms. MeSH descriptors are hierarchically organized into sixteen predefined groups, named top-level MeSH categories [5], that represent general topics such as anatomy, organisms, and diseases [6], so that each MeSH descriptor represents a narrower topic of at least one broader MeSH descriptor or MeSH category. MeSH descriptors are widely used in biomedical knowledge systems, primarily for the semantic indexing of scientific literature. In the MEDLINE part of PubMed [7], in particular, publications are annotated with topic labels primarily using MeSH descriptors. A process that used to be done by human indexers and has gradually transitioned to automation [8].

To build a comprehensive literature dataset we combined literature records with detailed affiliation information. This required matching publications that included both MeSH descriptors and author affiliations. Specifically, out of all the **916**,**824 publications initially retrieved from OpenAlex, 712**,**749 publications were annotated with MeSH descriptors and affiliation information, and out of these, 3**,**514 included at least one Greek affiliation**. These MeSH annotations, representing the main topics of each publication, were used in order to investigate which topics had higher or lower frequency in publications from Greece compared to publications from the rest of the world, as described in Methods. The data source for MeSH annotations was the 2024 version of the PubMed Baseline Repository [9], and MeSH annotations were available for about 78% of the publications, as PubMed has specific inclusion criteria related to the scope and quality of the publication journal [10].

### Thematic and statistical differences across MeSH categories

#### A. General overview Greece vs world (High-level MeSH Categories)

At the top level, comparing MeSH Categories, which are the roots of the MeSH polyhierarchy, we observed that the frequency of publications from Greece was higher in the “Diseases” category and lower in the categories of “Anthropology, Education, Sociology, and Social Phenomena” and “Psychiatry and Psychology”, which was statistically significant, as shown in **Table 1**. The remaining MeSH categories had smaller differences with estimate value below 0.0004, or were less significant (or not significant at all) as seen in **Table 1**. To note, the significance symbols are defined as follows (used consistently across the tables): p > 0.05 “ns” (not significant), p ≤ 0.05 “^*^”, p ≤ 0.01 “^**^”, p ≤ 0.001 “^***^”, and p ≤ 0.0001 “^****^”.

**Table 1:** Comparing the frequency of publications from Greece against publications from the rest of the world at the level of MeSH categories (Statistical analysis). Ranked by estimate.

| MeSH Category (level 0) | estimate | stat. sign. |
| --- | --- | --- |
| Diseases | 0.0006 | *** |
| Information Science | 0.0003 | ** |
| Chemicals and Drugs | 0.0001 | ns |
| Technology, Industry, and Agriculture | 0.0000 | ns |
| Health Care | 0.0000 | ns |
| Humanities | -0.0000 | ns |
| Analytical, Diagnostic and Therapeutic Techniques, and Equipment | -0.0000 | ns |
| Phenomena and Processes | -0.0001 | ns |
| Anatomy | -0.0001 | * |
| Geographicals | -0.0001 | ** |
| Disciplines and Occupations | -0.0002 | ns |
| Named Groups | -0.0002 | ** |
| Organisms | -0.0002 | ns |
| **Anthropology, Education, Sociology, and Social Phenomena** | **-0.0004** | **\*\*\*\*** |
| **Psychiatry and Psychology** | **-0.0007** | **\*\*\*\*** |

#### B. General overview Greece vs the world (mid-level MeSH categories)

Moving one level below, that is, to direct descendants of the MeSH Categories, we encounter 105 “MeSH subcategories” of level 1. Of them, only 17 had a notable difference in frequency of publications from Greece compared to the rest of the world (> 0.0001), which was also statistically significant (^****^, or ^***^), as shown in **Table 2**. For most of them (∼76%), the difference was negative, revealing some topics where the relative frequency of publications from Greece was found to be lower than the rest of the world. For four subcategories, however, the difference was negative, indicating that these topics are more frequent in publications from Greece than the rest of the world, proportionally.

**Table 2:** Comparing the frequency of publications from Greece against publications from the rest of the world at the level of MeSH subcategories (MeSH descriptors of level 1). Only cases that are notable (> 0.0001) and significant at level 1 are presented, ranked by estimate. The respective MeSH category is shown for completeness.

| MeSH Category (level 0) | MeSH Subcategory (level 1) | estimate | stat. sign. |
| --- | --- | --- | --- |
| Phenomena and Processes | **Mathematical Concepts** | 0.0003 | **** |
| Analytical, Diagnostic and ... | **Diagnosis** | 0.0003 | **** |
| **Diseases** | **Neoplasms** | 0.0002 | **** |
| **Diseases** | **Skin and Connective Tissue Diseases** | 0.0001 | **** |
| **Psychiatry and Psychology** | **Social Sciences** | -0.0001 | *** |
| Phenomena and Processes | **Biological Phenomena** | -0.0001 | **** |
| **Psychiatry and Psychology** | **Behavioral Disciplines and Activities** | -0.0001 | **** |
| **Anthropology, Education ...** | **Education** | -0.0001 | **** |
| Anatomy | **Nervous System** | -0.0001 | **** |
| Phenomena and Processes | **Chemical Phenomena** | -0.0001 | **** |
| Phenomena and Processes | **Musculoskeletal and Neural Physiological** | -0.0002 | **** |

|  | Phenomena |  |  |
| --- | --- | --- | --- |
| Phenomena and Processes | Genetic Phenomena | -0.0002 | **** |
| Phenomena and Processes | Physiological Phenomena | -0.0002 | **** |
| Health Care | Health Care Economics and Organizations | -0.0002 | **** |
| Health Care | Health Care Facilities, Manpower, and Services | -0.0002 | **** |
| Psychiatry and Psychology | Behavior and Behavior Mechanisms | -0.0003 | **** |
| Psychiatry and Psychology | Psychological Phenomena | -0.0003 | **** |

**Table 2** also reveals that differences observed at one hierarchical level are not necessarily distributed evenly into subordinate levels, where differences of varying magnitude and significance may be observed, even in the opposite direction. The MeSH subcategory “Mathematical Concepts”, for instance, has a positive difference, which is also significant, despite being included in the broader category “Phenomena and Processes” that has a slightly negative difference, overall, which is not substantial. At the same time, the “Phenomena and Processes” category includes four subcategories with a notable and significant negative difference. A similar situation holds for the “Diagnosis” subcategory, which presents a notable positive difference in frequency, despite belonging to a Category without a substantial difference overall. The category “Health Care”, on the other hand, with an insignificant positive difference overall, includes two subcategories with a significant negative difference.

#### C. Selected cases in comparison

Moving to descriptors at level 2 of the MeSH hierarchy, that is, direct descendants of MeSH subcategories, we can observe that an overall frequency difference that is notable and significant in a higher-level category is often explained by respective differences in only a few descriptors at a lower level. **Table 3**, for instance, reveals that the lower frequency of the “Psychiatry and Psychology” category in Greece is mainly explained by substantially lower frequencies in just nine level-2 descriptors. The remaining fifty level-2 descriptors that belong to “Psychiatry and Psychology” present only insignificant and/or too small differences (< 0.0001).

**Table 3:** Comparing the frequency of publications from Greece against publications from the rest of the world, considering MeSH descriptors of level 2 that belong to MeSH Category “Psychiatry and Psychology”. Only selected cases that are notable (> 0.0001) and significant at level 2 are presented, grouped by Subcategory, and ranked by estimate.

| MeSH Category (level 0) | MeSH Subcategory (level 1) | MeSH descriptor (level 2) | estimate | stat. sign. |
| --- | --- | --- | --- | --- |
| Psychiatry and Psychology | Behavior and Behavior Mechanisms | Behavior | -0.0002 | **** |
|  |  | Psychology, Social | -0.0001 | **** |
|  |  | Attitude | -0.0001 | *** |
|  |  | ... | ... | ... |
|  | Behavioral Disciplines and Activities | Psychophysiology | -0.0001 | **** |
|  |  | Sociology | -0.0001 | **** |
|  |  | Psychology, Social | -0.0001 | **** |
|  |  | ... | ... | ... |
|  | Psychological Phenomena | Mental Processes | -0.0002 | **** |
|  |  | Psychophysiology | -0.0001 | **** |
|  |  | ... | ... | ... |
|  | Social Sciences | Sociology | -0.0001 | **** |
|  |  | ... | ... | ... |
| ... | ... | ... | ... | ... |

Similarly, the positive difference in the “Diseases” category is mainly explained by respective positive differences in just four level-2 descriptors, as shown in **Table 4**, with the tens of their remaining sibling descriptors at the same level presenting insignificant frequency differences between Greece and the rest of the world. In addition, only two of these four level-2 descriptors belong to a subcategory (level 1) with notable and significant positive difference.

**Table 4:**
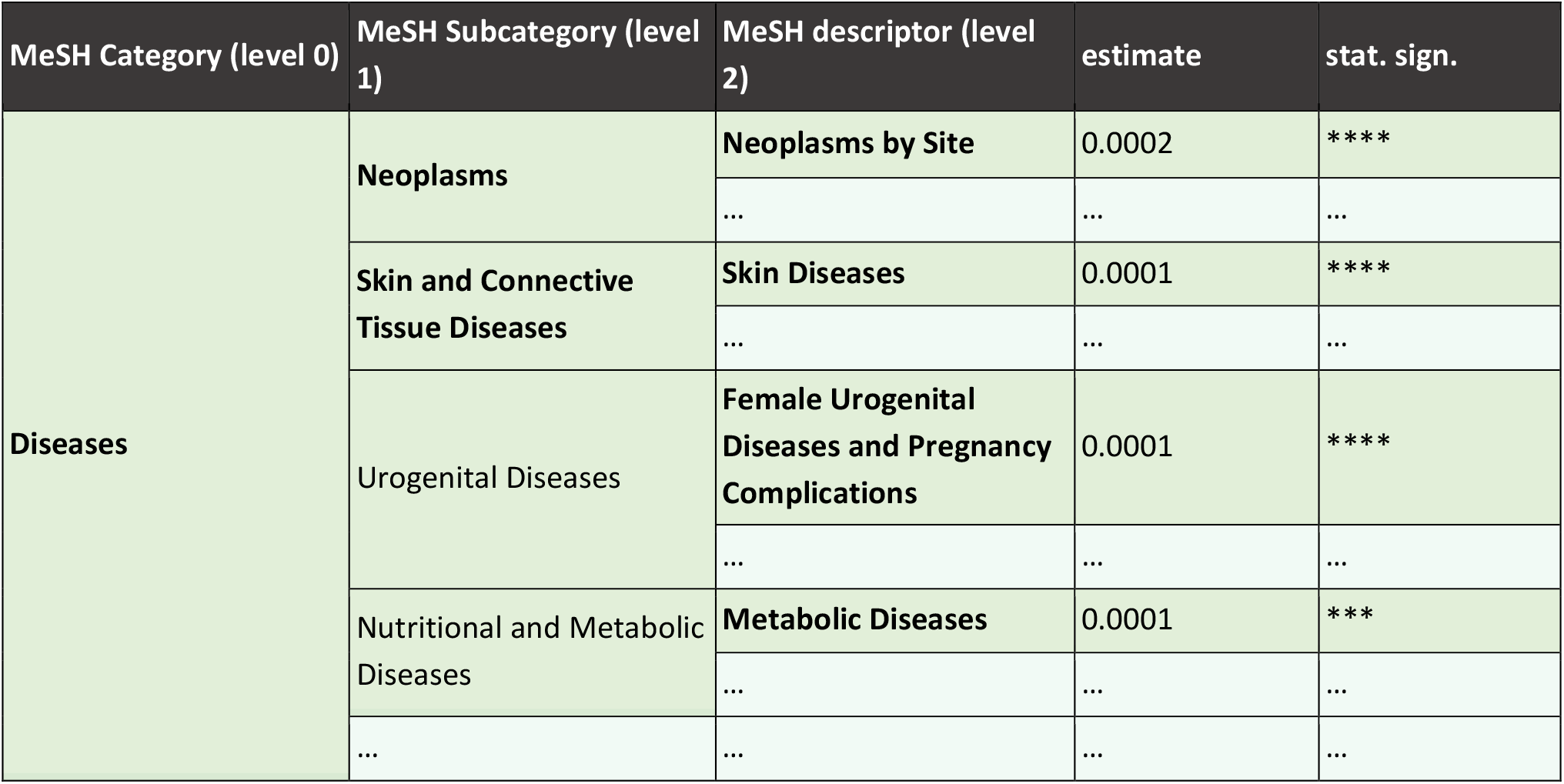
Comparing the frequency of publications from Greece against publications from the rest of the world, considering MeSH descriptors of level 2 that belong to MeSH Category “Diseases”. Only selected cases that are significant at level 2 are presented, grouped by Subcategory, and ranked by estimate.

Finally, considering this level of the polyhierarchy, the phenomenon of conflicting frequency differences inside the same category becomes even clearer. In particular, this happens in two categories, namely “Phenomena and Processes” and “Analytical, Diagnostic and Therapeutic Techniques, and Equipment”, as shown in **Table 5**. In addition, the level-2 descriptor “Physical Stimulation”, despite presenting a notable and significant difference, belongs to the subcategory “Investigative Techniques” that does not. This may suggest that a smaller frequency increase in several other level-2 descriptors inside this subcategory may counteract the effect of the decreased frequency of publications for “Physical Stimulation”.

**Table 5:**
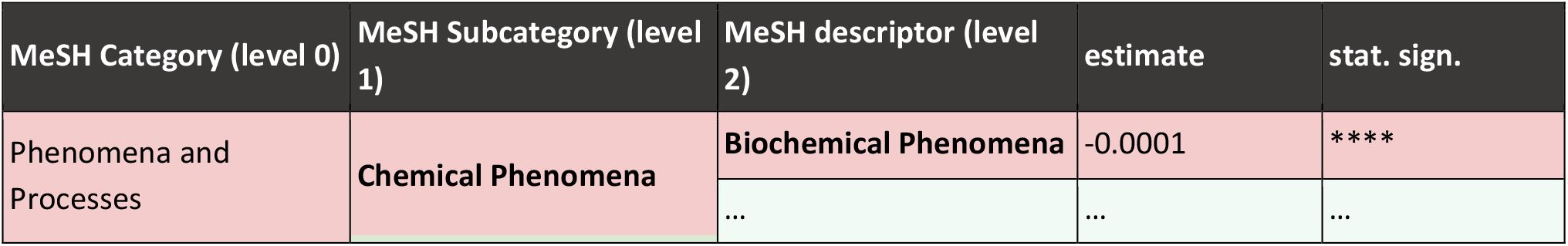
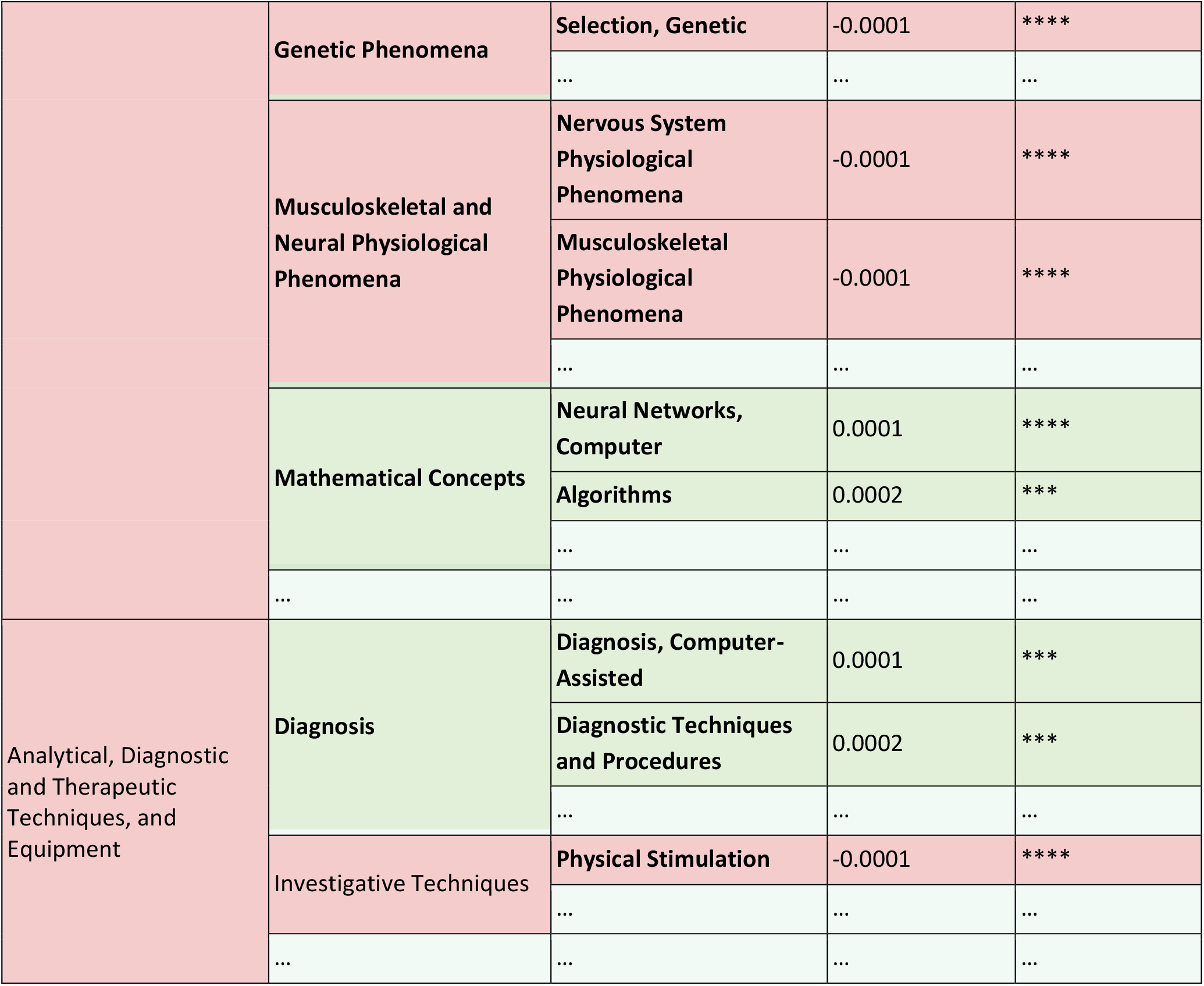
Comparing the frequency of publications from Greece against publications from the rest of the world, considering MeSH descriptors of level 2 that belong to MeSH Categories “Phenomena and Processes” and “Analytical, Diagnostic and Therapeutic Techniques, and Equipment”. Only selected cases that are significant at level 2 are presented, grouped by Subcategory, and ranked by estimate.

### Community perspectives from the national survey

As part of our effort to capture community perspectives on AI activity in the Life Sciences in Greece, we conducted a small but targeted survey leveraging established national networks in Greece including ELIXIR-GR, HSCBB and HBio. The survey questions and the respective anonymized responses used in the analysis below can be found on ELIXIR-GR Zenodo repository [11,12]. The overall participation was lower than anticipated (35 responses across 10 organizations); however, some indications could be derived from the captured information.

The survey results offer an overview of Greek AI–Life Sciences participants and their disciplinary backgrounds. The distribution of expertise (see **Figure 1**) shows a clear concentration in computational and analytical fields, including bioinformatics, computational biology, computer science, and software development.

**Figure 1:**
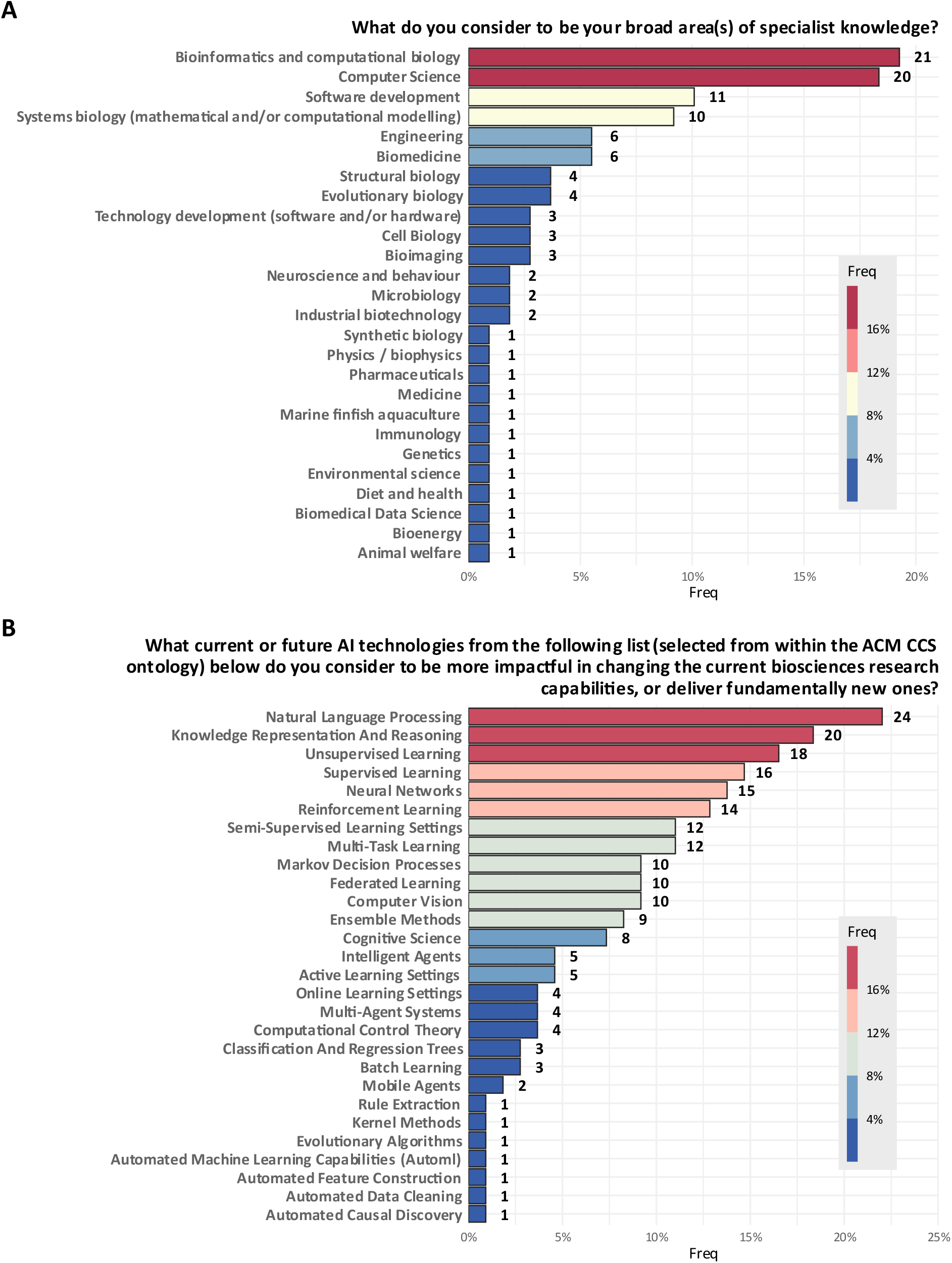
**(A)** Bar plot of the responses to the question #4 “What do you consider to be your broad area(s) of specialist knowledge?”. **(B)** Bar plot of the responses to the question #13 “What current or future AI technologies from the following list (selected from within the ACM CCS ontology) below do you consider to be more impactful in changing the current biosciences research capabilities, or deliver fundamentally new ones?”.

## Discussion

The findings of the systematic analysis of the literature highlight a distinct thematic divergence between Greece and global trends in the use of AI techniques within the Life Sciences. In Greece, the focus appears to be on the foundational development of tools, methods, and machine learning algorithms, likely driven by computer science and engineering departments. By contrast, international efforts tend to be led by life sciences departments, where the main driver is basic **biomedical research**. This divide shapes the nature of AI-related work and priorities across each context.

A closer look at selected MeSH subcategories further illustrates these differences in research orientation. or example, “Mathematical Concepts” and “Diagnosis” appear with higher relative frequency in Greek-affiliated publications, pointing to a strong national footprint in the development of computational methodologies and diagnostic tools. Conversely, topics such as “Behavior and Behavior Mechanisms” and “Psychological Phenomena” are markedly underrepresented compared to the global corpus, suggesting a lower degree of engagement with clinically embedded or patient-centered AI applications. Interestingly, within the broader “Diseases” category, Greek research shows a relatively higher focus on “Neoplasms” and “Skin and Connective Tissue Diseases,” whereas internationally the emphasis is distributed more evenly across multiple disease areas, including cardiovascular and neurological disorders. This pattern indicates that national strengths align with core algorithmic development and selected biomedical domains, while translational and behavioral health applications remain comparatively underexplored.

These bibliometric insights resonate with the findings of the national survey, which, although limited in scale (∼35 responses), provides valuable community perspectives on why such gaps persist in practice. When asked about the key challenges in delivering AI-related research goals in Life Sciences, respondents consistently pointed to three major areas: data issues, infrastructure and resources, talent and collaboration. Data-related issues include poor quality, scarcity, lack of standardization and interoperability, and difficulties in accessing real-world datasets due to regulatory and proprietary constraints. Infrastructure challenges involve limited computational resources, especially GPU access, and constrained funding. Additionally, there is a shortage of skilled personnel and insufficient collaboration between AI and life sciences disciplines.

In terms of required competencies, the survey responses indicated a strong emphasis on foundational expertise in mathematics, statistics, and computer science. Basic knowledge of linear algebra, algorithms, computational statistics, and systems programming are in high demand but hard to find in combination with deep cross-disciplinary understanding. It is clear that there is a significant need for professionals who can integrate AI capabilities with domain-specific biological insights and apply them meaningfully. Furthermore, respondents emphasized the difficulty in recruiting software engineers capable of robust implementation and adaptation to evolving AI architectures.

To promote stronger community-level engagement in AI bioscience in Greece, participants suggested a variety of activities. These include educational and training events such as workshops, seminars, summer schools, and Master’s programs that offer real-life project experience and certification. Collaborative forums like hackathons, datathons, and online discussion platforms were also recommended to foster interdisciplinary exchange. Additionally, involving startups, hospitals, and the public through outreach, awareness campaigns, and industry collaborations was seen as vital to building momentum.

The proposed activities are expected to bring several key benefits. First, they would enhance collaboration and networking, particularly across disciplines and career stages, benefiting students and early-career researchers. Second, they would support skill and knowledge development by strengthening computational expertise among biomedical scientists and increasing domain understanding among AI specialists. Finally, these efforts would help build trust and awareness, improving public and scientific confidence in AI technologies and supporting more informed and ethical integration into everyday practice.

It should be noted that, based on the overview of the available MSc programs, this perspective is also evident, as overall, the identified programs aim to equip students with interdisciplinary skills combining data science, informatics, and engineering to address complex challenges in biomedical research, healthcare innovation, and translational medicine.

Finally, it is worth mentioning that the findings of this work are reinforced by independent policy documents. A Key example is the 2022 Policy Support Facility (PSF) Final Review report to Greece for policies developing research infrastructures and the R&I ecosystem [13], in which it is stated that “[…*] Greece also has a very strong background in informatics and computer science and given the growing visibility of artificial intelligence (and the importance this has in the Greek strategic planning)[*…]”, as well as the fact that *AI can blend well with bioinformatics applications*. Both of these points reinforce the findings of our analysis in the unique strengths of the Greek scientific community in the use of AI techniques within the Life Sciences.

## Methods

Data retrieval, annotation, and filtering were carried out in successive stages, followed by a systematic statistical analysis between the selected groups. All code that has been developed specifically for this analysis, is openly and publicly available under an MIT license in the GitHub repository https://github.com/npechl/Elixir-GR-ML.

### Data sources and retrieval

OpenAlex served as the principal source for bibliographic information for this study. As one of the most comprehensive Scholarly Knowledge Graphs (SKGs), OpenAlex provides extensive metadata for research products (e.g. publication and datasets) as well as their relationship with other research-related entities (e.g., organizations, scientific topics).

In this study, we focused on publications explicitly linked with the OpenAlex top-level concepts of “**Medicine**” and “**Biology**”, and at the same time categorized under “**Natural Language Processing**”, “**Artificial Intelligence**”, or “**Machine Learning**”. or each retrieved publication, we collected rich metadata including title, authors, affiliations, DOI, PMID, and topic annotations (the first two levels of OpenAlex concepts hierarchy).

The entire data acquisition process was implemented using Python, leveraging the October 2024 OpenAlex snapshot [14] to retrieve, filter and transform relevant metadata into a convenient format for further analysis.

Subsequently, each DOI was linked with its corresponding Medical Subject Heading (MeSH) terms as cataloged in PubMed. MeSH terms constitute a regulated vocabulary employed by the National Library of Medicine for the indexing and organization of biological content. They are arranged in a hierarchical framework comprising many levels, from broad categories (e.g., “Diseases” or “Analytical, Diagnostic, and Therapeutic Techniques”) to more specific, detailed descriptions.

All data are available at https://doi.org/10.5281/zenodo.17415352 and https://doi.org/10.5281/zenodo.17407809.

### Data curation (pre-processing)

Multiple preprocessing steps were implemented to ensure the robustness of the study, all carried out using R (R-4.4.0). The dataset was divided into two categories:articles associated with Greek institutions (group “Greece”) and all other publications (group “World”). A publication was assigned to the “Greece” group if it listed at least one institution located in Greece.

Next, we calculated the frequency of each MeSH term for both groups over the period 2000 – 2020. Since MeSH terms are organized hierarchically across multiple levels; this study focused on terms from the first three levels. To mitigate potential bias due to differences in annual publication volumes, relative abundances were calculated by normalizing MeSH term frequencies against the total number of publications for each year. Finally, z-scoring was applied where appropriate (i.e., in heatmaps) to highlight the differences between the two groups.

All code used for this preprocessing is available at https://github.com/npechl/Elixir-GR-ML.

### Statistical analysis

To identify topics with significant differences between the two groups, Wilcoxon signed-rank tests [15] were performed for each year, comparing the relative abundances of individual MeSH terms. P-values were adjusted using the Holm method [16] to account for multiple testing. The results were then compiled into a color-coded, multi-level table, linking each level 2 topic to its corresponding level 1 and level 0 ones. This allowed us to identify patterns across the MeSH hierarchy.

The final step aimed to identify topics exhibiting notable differences between Greece and the rest of the world. Specifically, we focused on topics with an estimated difference – calculated as the median of the differences between samples from “Greece” and “World” – greater than 10^−5^. To balance term generality and specificity, while reducing potential noise, we restricted our analysis to topics classified up to the second level of the MeSH term hierarchy.

### Community Assessment

#### A survey on Machine Learning for Life Sciences in Greece

A complementary national survey, modeled on a similar UK study [1] was designed and distributed to research teams across the Greek scientific community to gather subjective insights into the current landscape. Implemented via Google Forms, the survey explored both the scientific domains and research practices of these teams.

It comprised 18 questions, covering a range of topics: from general descriptive information (e.g., age, disability status, group size, scientific background, and research specialization) to more AI-focused aspects, such as the extent of AI/ML involvement in their work, methodologies and tools used, and their views on the broader community’s engagement with AI/ML. The full survey structure and questions can be found here.

#### Mapping of training programs in Greece

Additionally, we performed a review of available courses (under-/post-graduate level) in Greece on the AI/LS intersection. To this end, we compiled and shared a list of master’s degree programs offered in Greece that focus on the intersection of machine learning or artificial intelligence and biology. We collected 12 courses from 8 public universities and research organizations in Greece offering in person, online or hybrid master courses.

These postgraduate programmes focus on advanced topics at the intersection of data science, bioinformatics, biomedical engineering, medical informatics, and AI. A systematic overview of the provided descriptions and available content across these programmes, showed that the key themes include:

- Data science & big data analytics applied to health, life sciences, and bioinformatics.
- Bioinformatics and computational biology, including NGS data analysis and use of AI technologies (e.g., machine learning, deep learning, neural networks, knowledge graphs).
- Precision and personalized medicine, emphasizing molecular and genomic data for individualized therapy.
- Biomedical engineering, covering areas such as biofabrication, molecular diagnostics, imaging, and medical information systems to support precision medicine.
- Health statistics and data analytics, training graduates to analyze and interpret health data for research, public health, and clinical applications.
- Digital health and AI in healthcare, focusing on the development and use of advanced digital tools, ICT, and AI technologies in health systems.

It should be noted that some programmes also include modules on computational medicine, simulation, computer security, and the application of ICT in education and healthcare.

## Software and Data availability

Publications related to AI, ML, and NLP within Medicine and Biology (derived from OpenAlex) that were used for the present study are available at https://doi.org/10.5281/zenodo.17415352. The survey on Machine Learning for Life Sciences in Greece along with the anonymized responses can be found at https://doi.org/10.5281/zenodo.17287162 and https://doi.org/10.5281/zenodo.17414392.

All R scripts used in this study and any other analysis files are available at https://github.com/npechl/Elixir-GR-ML.

## Acknowledgements

This work was supported by ELIXIR, the research infrastructure for life science data.

## Author contributions statement

A.D., N.P. and A.N. analyzed the data and produced the tables and the figure. S.C. and K.V. gathered and processed the data from OpenAlex. All authors contributed to the writing and the review of the manuscript.

## Additional Information

The authors declare no competing interests.

